# Transmission risk assessment of *Aedes*-borne arboviral diseases in Accra, Ghana

**DOI:** 10.1101/2022.02.14.480316

**Authors:** Nukunu E. Akyea-Bobi, Jewelna Akorli, Samuel Sowah Akporh, Dominic Acquah-Baidoo, Millicent Opoku, Kwadwo Frempong, Sellase Pi-Bansa, Helena A. Boakye, Joannitta Joannides, Mufeez Abudu, Joseph Harold Nyarko Osei, Rebecca Pwalia, Esinam Abla Akorli, Godwin K. Amlalo, Kofi Bonney, Reginald Quansah, Samuel K. Dadzie

**Affiliations:** Vestergaard NMIMR Vector Labs, Parasitology Department, Noguchi Memorial Institute for Medical Research, University of Ghana, P.O. Box LG 581, Legon, Accra; Department of Parasitology, Noguchi Memorial Institute for Medical Research, University of Ghana, P.O. Box LG 581, Legon, Accra; Department of Virology, Noguchi Memorial Institute for Medical Research, University of Ghana, P.O. Box LG 581, Legon, Accra; Department of Biological, Environmental and Occupational Health, School of Public Health, University of Ghana, P.O. Box LG 13, Legon, Accra

## Abstract

**Background:** Dengue, Zika and Chikungunya are *Aedes-borne* viral diseases that have risen to become great global health concerns in the past years. Several countries in Africa have reported outbreaks of these diseases and despite Ghana sharing borders with some of such countries, it remains free of these outbreaks.

Recent studies in Ghana have revealed that there are antibodies and viral RNA of the Dengue virus serotype-2 among individuals in some localities in the Greater Accra Region. This is an indication of a possible silent transmission ongoing in the population, hence the need to assess the risk of transmission of these viruses within the country. This cross-sectional study, therefore, assessed the risk of transmission of Dengue, Zika and Chikungunya viruses in a domestic/peri-domestic (Madina) and a forest (Achimota Forest) population in the Greater Accra Region, Ghana.

**Methodology/Findings:** All stages of the *Aedes* mosquito (egg, larvae, pupae and adults) were collected around homes and in the forest area for estimation of risk indices. All eggs and immature stages were reared to adults and morphologically identified. The predominant species of *Aedes* mosquitoes identified from both sites were *Aedes aegypti* (98 % in Madina and 98.1% in Achimota forest). *Aedes albopictus*, an important arbovirus vector, was identified only in Madina at a prevalence of 1.5% but Achimota forest had the higher species diversity. Both study sites recorded high risk indices; Madina: Positive Ovitrap Index = 26.6%, Container Index = 36.8%, House Index = 19.8%, Breteau Index = 70.4%; Achimota: Positive Ovitrap Index = 34.2% and Container Index = 67.9%. RT-PCR to detect the presence of Dengue, Chikungunya and Zika viruses was negative for all pools tested.

**Conclusion:** All entomological risk indicators estimated showed that both sites had a high potential of an outbreak of arboviral diseases following the introduction of these viruses.

**Author Summary:** The detection of antibodies and viral RNA of the dengue virus serotype 2 in some communities in the urban city of Accra, suggested the possibility of silent transmission of arboviral disease within the city. We assessed the risk of arboviral disease transmission using entomological risk indices. The study was a cross-sectional study conducted in a forest and peri domestic setting located in the southern urban city of Accra.

The different stages of the *Aedes* mosquito were collected and, houses and containers positive for *Aedes* mosquitoes were also noted. The Breteau (BI), House (HI), Container (CI) and Positive ovitrap (POI) indices were determined. Real Time-PCR was conducted to determine the presence of Dengue, Zika and Chikungunya viruses in the larvae and adults collected.

*Aedes aegypti* was the most common species identified from both sites. *Aedes albopictus* another competent arbovirus vector was identified in the peri-domestic site. Almost all risk indices recorded for both sites were higher than the WHO thresholds allowed for these indices. However, real time-PCR to detect the presence of Dengue, Chikungunya and Zika viruses was negative.

The high entomological risk indicators estimated showed that both sites had a great potential of an outbreak following the introduction of these viruses, and a well-structured surveillance for these vectors is highly recommended. The detection of the presence of *Ae. albopictus*, an invasive species is also of great concern.

## Introduction

Dengue, Zika and Chikungunya viruses are all transmitted in urban and peri-urban areas by *Aedes* mosquitoes of the subgenus *Stegomyia*. All three viruses are transmitted in zoonotic cycles involving primates and mosquitoes (1). However, human activities such as the domestication of the natural habitats of the reservoirs and vectors of these viruses have led to an urban transmission cycle involving human-to-human transmission where *Aedes aegypti* and *Aedes albopictus* are major vectors (2,3).

*Aedes* mosquitoes are highly invasive species found on almost all continents. It has one of the widest distributions ever recorded (4). *Aedes aegypti* is mainly found in the tropics and subtropics, flourishing in domesticated environments and has adapted to feeding almost solely on humans. It is active during the day and usually bites several people during the acquisition of a bloodmeal. These behavioural characteristics of this mosquito combined with its high susceptibility to the Dengue, Zika and Chikungunya viruses make it a very efficient vector (5).

Ecological and human factors seem to be strong determinants for the spread of arboviruses and their vectors. The biology of the vectors, their abundance, and geographical distribution are directly affected by climate (6). Global trade such as the importation of used tyres, ornamental plants and used vehicles is also one of the driving factors for the spread of *Aedes* mosquitoes (7). Infected travellers also enable the virus to reach new places within the shortest possible time (8). This can lead to an explosive epidemic in areas where there is a well-established vector population. Other human activities such as poor planning of urban centres leading to issues like lack of consistent water supply, poor drainage and waste disposal systems also facilitate the development of vectors (9); most outbreaks of arboviral diseases in Africa have occurred in urban cities (10). Rapid urbanization has also been documented as one of the drivers of the spread of arboviral diseases (11).

In Africa, thirty-four (34) countries have reported Dengue with their capital cities being the hardest hit (12). Ghana shares borders with countries (Burkina Faso, Cote d’Ivoire and Togo) that have reported DENV, CHKV and ZIKV. However, Ghana has remained free of an outbreak of these diseases despite *Aedes aegypti* mosquitoes being present in most part of the country (13–15). Recent studies have revealed evidence of DENV-2 (Dengue virus serotype-2) antibodies in people suspected to have malaria or visiting the hospital with acute fever symptoms (16,17). The DENV-2 viral RNA was also detected among two children in some localities in urban Accra. Further inquiries from the parents of the infected children revealed that these children had never been out of the country before. This indicates that the infections were acquired locally, suggesting the virus is silently circulating in the population and there may be ongoing transmission (18). This study was aimed at determining the species composition, abundance, and distribution of *Aedes* mosquitoes and estimating the risk of transmission of arboviruses within a forest and some peri-urban communities in Accra, Ghana.

## Materials and methods

### Study area

The study was carried out in two sites: Achimota Eco-park (urban forest area) and Madina (peri-urban community) located in the capital city of Accra Metropolis, Ghana (Fig 1). Achimota is an eco-tourist site with easy access to potential visitors. The forest hosts the Accra zoo and the people that frequent this eco-park originate from various parts of the country. The presence of these patrons in the forest area where there are animals that serve as a reservoir of these viruses makes it a probable site for DENV, ZIKV and CHKV virus circulation.

**Fig 1:**
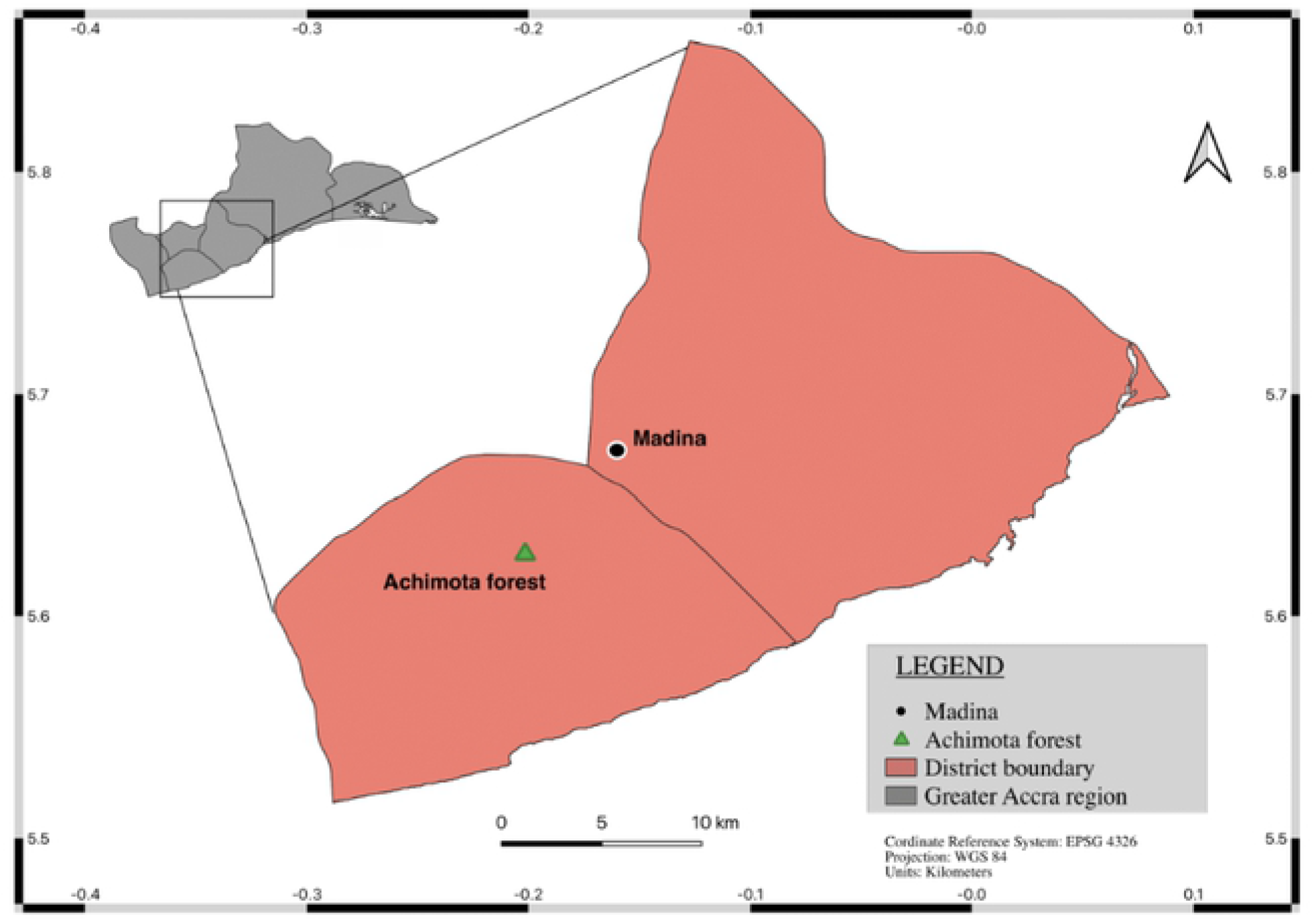
Map of Greater Accra region showing point locations of Achimota Eco-park and Madina.

Madina is an urban area which has over several years developed into a central business district becoming a hub for major commercial activities. This has led to the influx of rural-urban migrants looking for better prospects. Mechanic shops and tyre sale shops are common businesses found within this community. These car spare parts and tyres found in these shops are imported from all over the world. Due to vertical transmission of these viruses, eggs attached to these substrates could potentially be carrying DENV, ZIKV and CHIKV (19). There is therefore the possibility of introducing *Aedes* mosquitoes carrying DENV, CHKV and ZIKV into the community.

### Mosquito Collection

The Geographical Positioning System (GPS) was recorded for each point within the study area where the various stages of the *Aedes* mosquito were sampled using the KoBoCollect and GPS Essentials app; the data was spatially displayed on a map. All collections were done within the months of July-September 2020.

#### Egg sampling

A cross-sectional entomological study was carried out in the Achimota forest and Madina Municipality. Ovitraps to collect *Aedes* eggs were prepared from plastic water bottles cut into two halves and painted black on the outside. Each trap was lined with brown paper and filled with approximately 300mL of hay-infused water. Each study site was divided into four quadrants and 10 ovitraps were placed in randomly selected points per quadrant. The ovitraps were left for five days to collect mosquito eggs. The traps were retrieved and replaced over three different time points within the months of July-September 2020. During retrieval, the brown paper lining was checked for the presence of eggs, removed from the trap and stored in labelled Ziploc bags. The hay-infused water was also checked for larvae, and if any, was poured into collection bags and returned to the lab. All samples were brought back to the Noguchi Memorial Institute for Medical Research for further analyses. The number of eggs per trap was counted with the aid of a microscope. This data was used for the estimation of the Positive Ovitrap Index (POI) and the Egg Density Index (EDI). Brown paper containing eggs and the water contents of the ovitraps were poured into larval trays in the laboratory and allowed to hatch. The emerging adults were identified using identification keys (20).

#### Larval and pupal sampling

Potential breeding sites in and around human dwellings in the study sites were scouted for *Aedes* larvae and pupae. The larvae and pupae collected were transported to the laboratory and counted for estimations of larval risk indices; House index (HI), Container Index (CI) and Breteau Index (BI).

#### Adult sampling

The Biogents-sentinel traps (BG-traps) were used to capture adult *Aedes* mosquitoes outdoors. Four (4) BG-traps were placed outside randomly selected houses and properties within a chosen community in each study area. Each study site was divided into four quadrants and one BG trap was placed per quadrant. The collection was done over a period of 12 hours (6:00 am to 6:00 pm). The adult mosquitoes caught by the trap were collected into sealed vials, transported to the laboratory and stored at −80°C for further analyses.

### Morphological Identification of *Aedes* mosquitoes

The adult and larval stages of the *Aedes* mosquitoes collected from the field were morphologically identified in the NMIMR entomology laboratory using morphological identification keys (Rueda, 2004). Identification was done with the aid of a stereomicroscope at 4x and 2x magnifications. After identification, the mosquitoes were grouped into pools of 20 for the larvae and groups of 10 for the adults according to species. All samples were preserved in RNALater (SIGMA^®^) at −80°C, for viral detection.

### Viral detection using Trioplex Real-time RT-PCR Assay

Each sample was homogenized in Minimum Essential Medium (MEM) containing Earle’s Salts, 2% L-glutamine, supplemented with 10% heat inactivated FBS and Penicillin-Streptomycin. The Qiagen Viral RNA mini Kit was used according to the manufacturer’s instructions to extract RNA from pools of mosquitoes. The AgPath-ID RT-PCR (Real-Time Reverse Transcriptase Polymerase Chain Reaction) kit was used to detect DENV, CHIKV and ZIKV according to the protocol by CDC (2017).

### Data analyses

The abundance of mosquitoes and proportions of different species of *Aedes* mosquitoes were calculated per site and compared using a Mann-Whitney test in Stata/IC 15.0. Three larval indices and two ovitrap indices were used to estimate the *Aedes* mosquito density in both study sites. The larval indices estimated were House index (HI-the percentage of houses infested with Aedes larvae or pupae), Container Index (CI-the percentage of containers infested with larvae or pupae) and Breteau index (BI-the number of positive containers per 100 inspected houses). The two larval indices estimated were positive ovitrap index (POI-percentage of ovitraps positive with *Aedes* eggs) and the egg density index (EDI – average number of eggs per positive trap). All indices estimated were compared to WHO thresholds for transmission risk of viral haemorrhagic fevers (VHFs) to determine the risk of transmission level per index.

## Results

### Spatial distribution of *Aedes* breeding sites

Maps were constructed with the recorded GPS coordinates of positive households and breeding sites in the study areas to show the spatial distribution of *Aedes* breeding sites within the study areas. Figure 2 & 3 shows the spatial distribution of positive breeding sites of *Aedes aegypti* larvae relative to human habitation in Achimota Forest and Madina.

**Fig 2:**
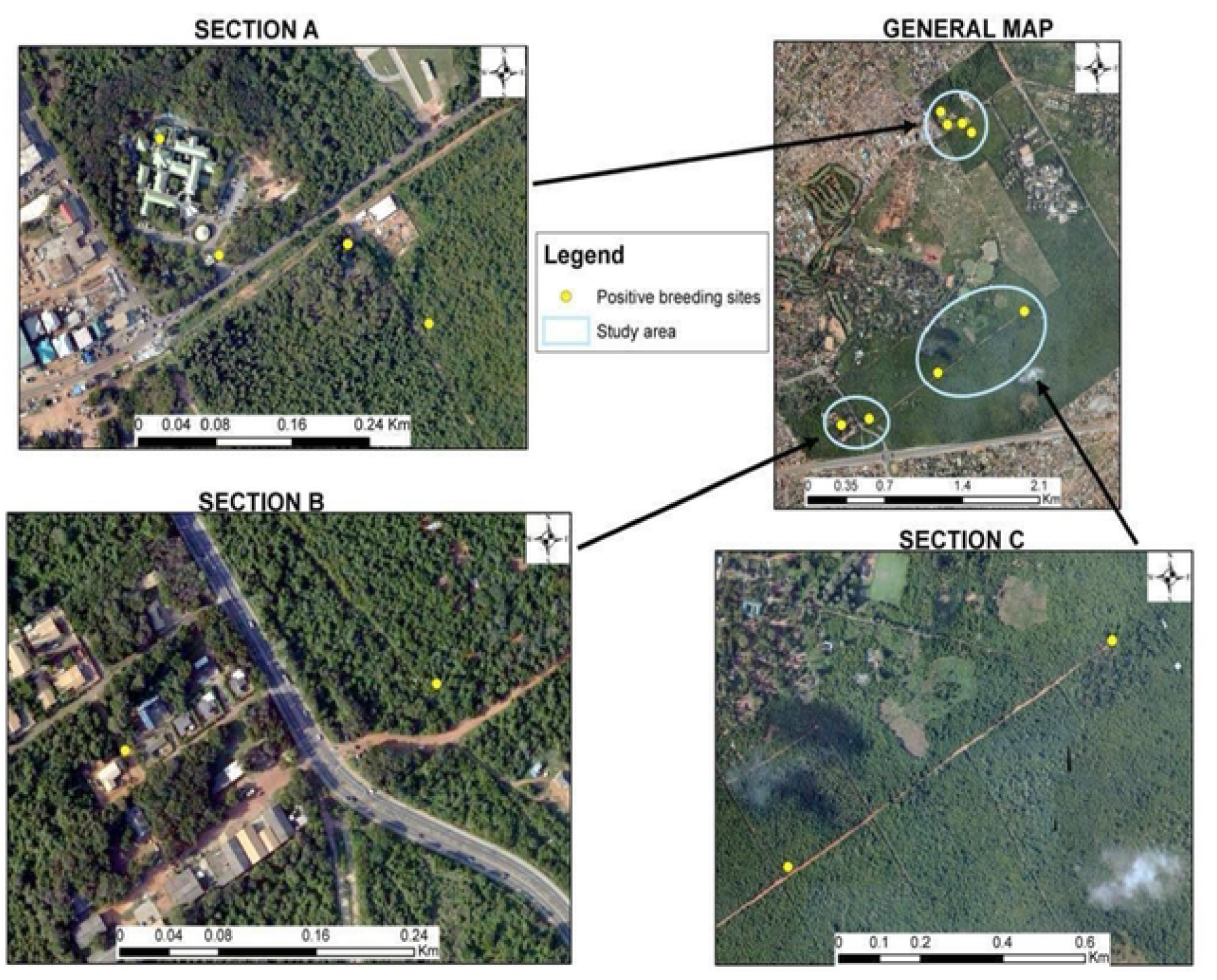
Spatial Map of Achimota eco-park showing positive breeding sites for Aedes larvae and pupae. The yellow-coloured dots show the sites of the containers found to be positive for Aedes larvae and pupae. Clustered sections on the map have been expanded to clearly show the distribution of the positive breeding sites.

**Fig 3:**
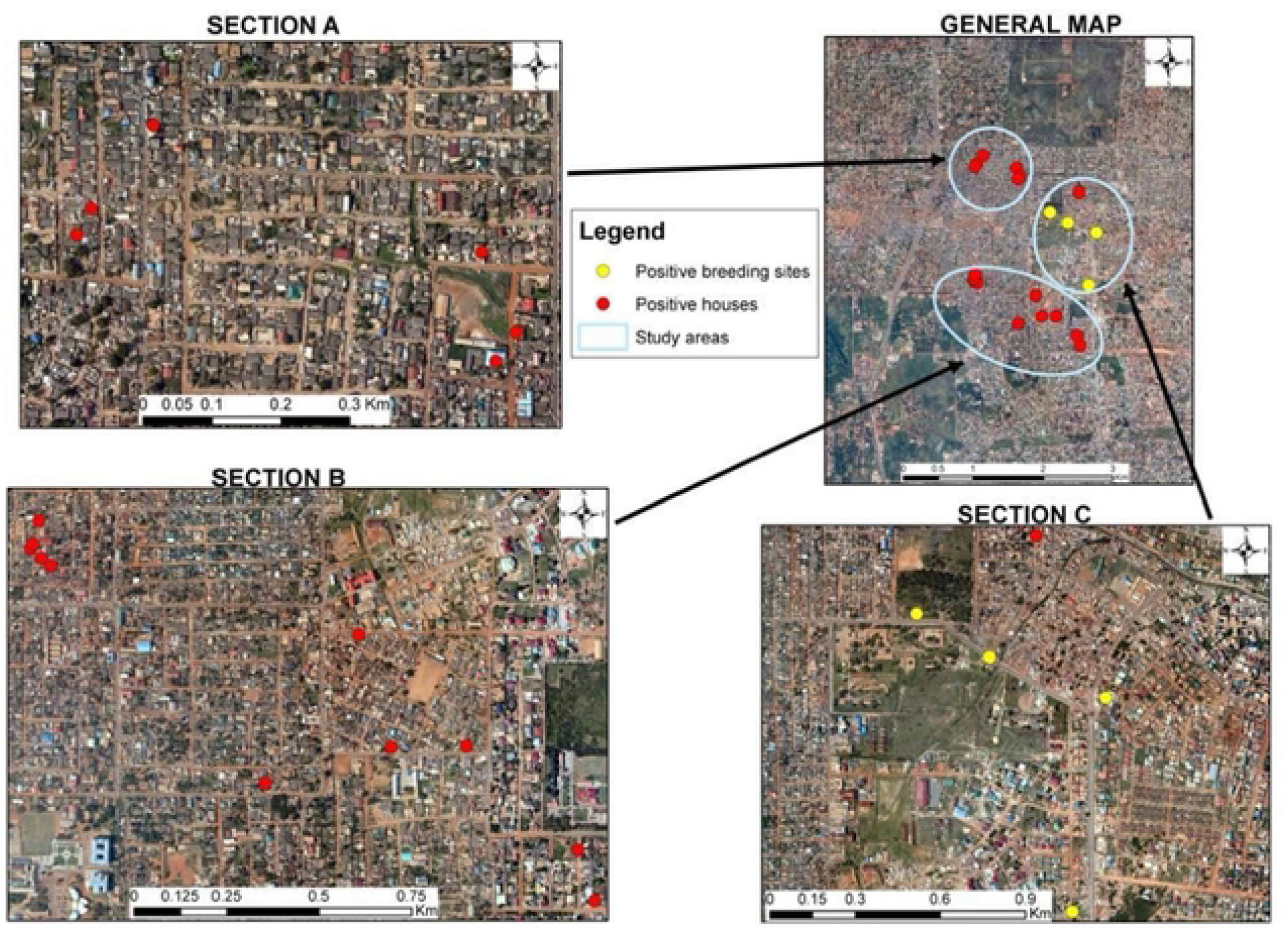
Spatial Map of Madina showing positive breeding sites for Aedes larvae and pupae. The yellow-coloured dots indicate positive containers outside the houses, and the red dots indicate houses with positive containers for *Aedes* larvae and mosquitoes. Clustered sections on the map were expanded to clearly show the distribution of the positive breeding sites.

### Species abundance and composition of *Aedes* mosquitoes

A total number of 120 ovitraps were set per study site. The total number of positive traps for Achimota forest and Madina were 38 and 29, respectively. The total number of eggs counted for Achimota forest and Madina was 1614 and 828, respectively. A total of 1150 larvae were collected from both sites; 560 for Achimota forest and 590 for Madina. The BG traps sampled a total of 8 adults from Achimota forest and 28 adult mosquitoes from Madina. There was no significant difference between the study areas in the number of samples collected (Fisher exact: *P*> 0.05)

A total of 2218 mosquitoes (adults and larvae) were identified to the species level (Fig 4). The most common species of *Aedes* mosquitoes identified from both sites were *Aedes aegypti;* 98% (1148) in Madina and 98.1% in Achimota (1027). Other species identified in Madina included *Aedes albopictus* (1.5%), *Aedes simpsoni* (0.2%) and *Aedes ingrami* (0.3%). Achimota forest recorded a higher number of *Aedes* species; these were *Aedes ingrami* (0.4%), *Aedes simpsoni* (0.3%), *Aedes domesticus* (0.4), *Aedes africanus* (0.7), *Aedes vittatus* (0.1%) and *Aedes de-boeri* (0.1%).

**Fig 4:**
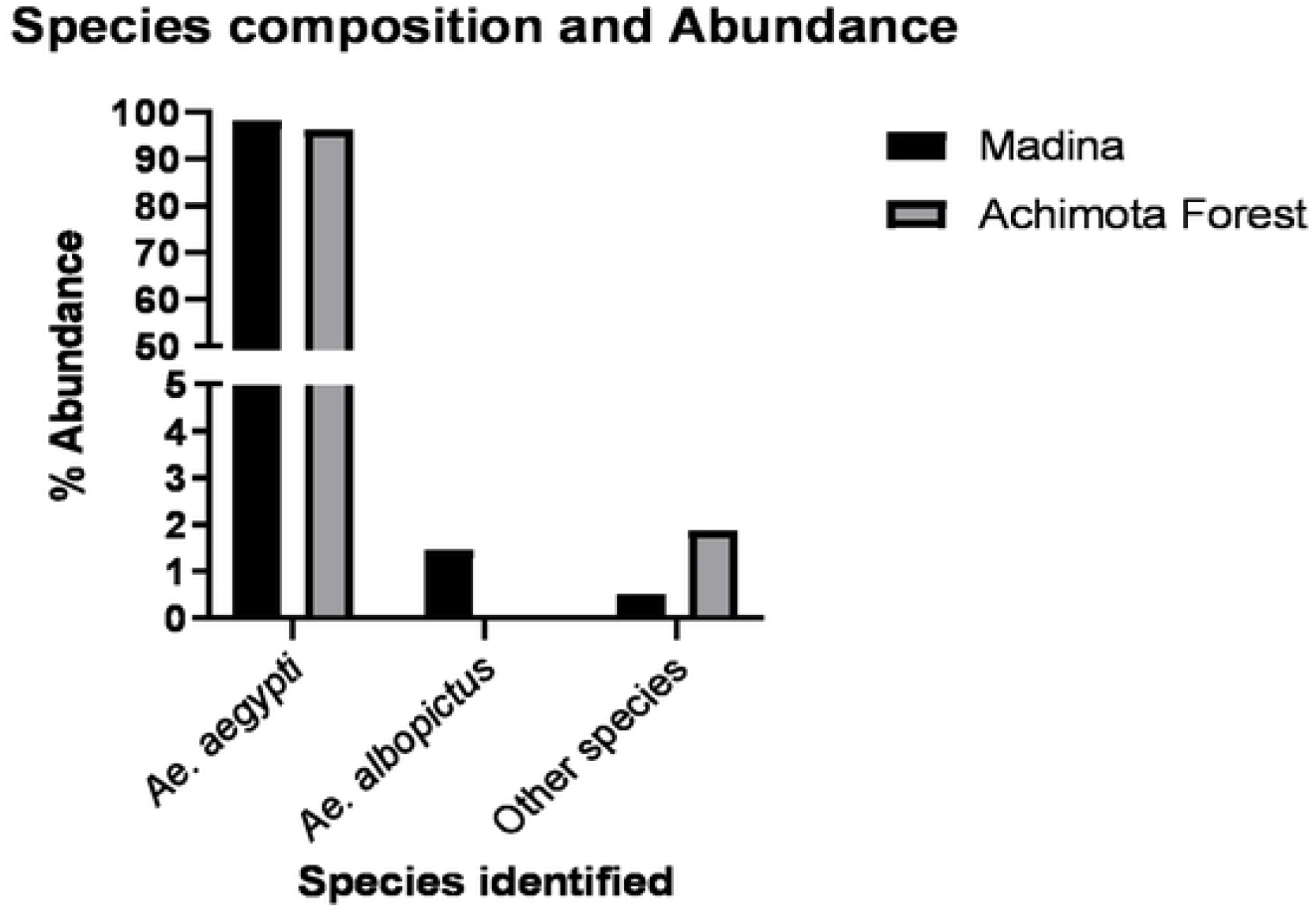
Species composition and abundance of *Aedes* mosquitoes identified from Achimota forest and Madina.

### Larval and Ovitrap indices estimation from household surveys

Larval and Ovitrap indices estimated in this study were above the WHO threshold except the House index in Madina which fell within the moderate risk range (Table 1). Out of 106 traps retrieved per site, Achimota forest and Madina had 38 (35.9%) and 29 (27.4%) ovitraps positive with *Aedes* eggs, respectively. This gave an egg density per trap of 29 in Madina and 43 in Achimota forest. During the house survey, a total of 81 houses were inspected in Madina and 16 (19.8%) were positive for *Aedes* larvae. No houses were surveyed in Achimota forest because there were no living quarters available. A total of 155 outdoor or peri-domestic containers were inspected in Madina; 57 (36.8%) were positive. In Achimota forest, 53 containers were inspected and 36 (67.9%) of these containers were found positive for *Aedes* larvae. Some examples of containers and breeding sites that were inspected included plastic and metal barrels, buckets, earthenware pots, Polytanks^®^ (large water storage tanks), abandoned bathtubs, abandoned machine parts and car tyres. Tyres were found to be the most positive containers for *Aedes* larvae and pupae in both sites (Fig 5).

**Table 1:** Larval and ovitrap indices for risk of transmission of Aedes-borne viral diseases in Madina and Achimota Forest

| Entomological indices for study areas | Madina | Achimota Forest | WHO Threshold for transmission risk of VHF's |
| --- | --- | --- | --- |
| Egg density index | 29 | 43 | NA |
| Positive ovitrap index | 27.4* | 35.5* | > 10 |
| Container index | 36.8* | 67.9* | 3-20 or above |
| House index | 19.8* | NA | 4-35 or above |
| Breteau index | 70.4* | NA | 5-50 or above |

(\*) indicates areas with high VHF transmission risk

**Fig 5:**
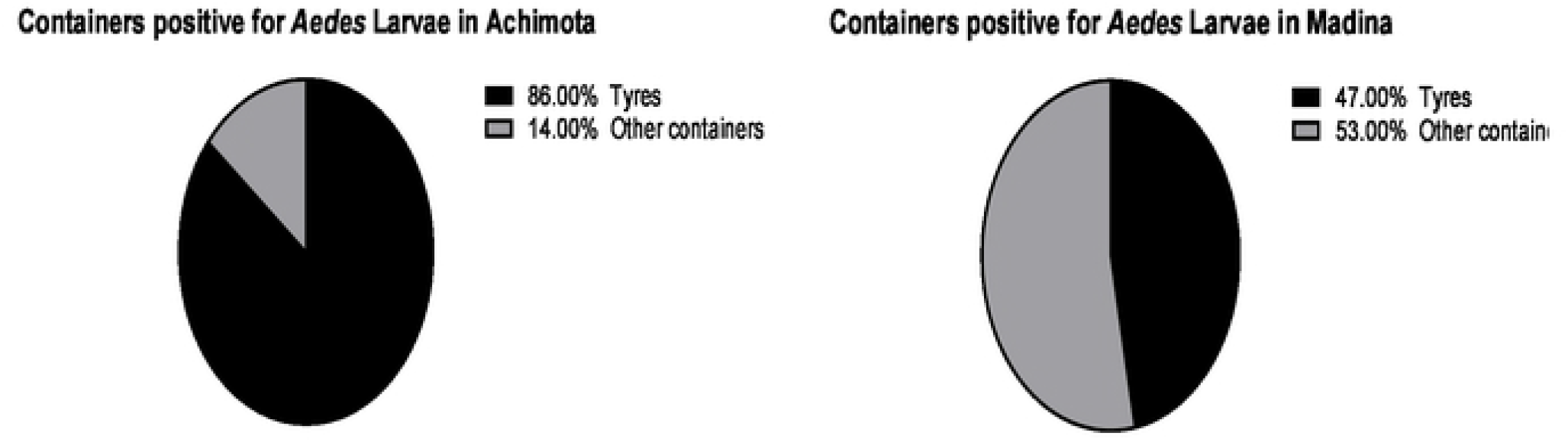
Distribution of containers positive for *Aedes* larvae and pupae in Madina and Achimota forest.

### Viral detection

A total of 120 mosquito pools was tested for the presence of DENV, ZIKV and CHKV. These consisted of a total of 1493 adult male and female *Aedes aegypti* mosquitoes and 1 pool of *Aedes albopictus* from both study sites totalling. All the pools tested negative for DENV, ZIKV and CHKV in both Achimota and Madina.

## Discussion

The presence of *Aedes* mosquitoes have been well documented in Ghana and have been implicated for the transmission of yellow fever (13,22). Studies have detected the presence of antibodies and viral RNA of DENV in different localities in Ghana (16,17). This has emphasized the need to investigate and understand the role of *Aedes* populations in the spread of these diseases in the country, which will inform the formulation of effective and efficient strategies to prevent future outbreaks or mitigate any outbreak that may arise.

This study focused on measuring the risk of transmission of *Aedes*-borne viral diseases, specifically DENV, ZIKV and CHKV in a domestic (Madina) area and in a forest area (Achimota forest). All stages of the mosquito were collected to detect their viral infectivity status. The most abundant species of the *Aedes* mosquito recorded in both populations was *Aedes aegypti*. This correlates with other studies in Ghana that have found *Aedes aegypti* to be the most predominant (15,23). The high abundance of these vectors in the country have been attributed to the presence of water holding containers close to human dwellings serving as breeding sites for these mosquitoes. *Aedes aegypti* has over the years evolved to live close to humans and preferentially choose to feed almost solely on humans even in the presence of other vertebrates that could serve as host (24). Therefore, the availability of breeding sites close to human dwellings may lead to a rise in the population of these *Aedes* species. From the spatial maps obtained for the sites, we observed the proximity of positive breeding sites to houses in the area; providing evidence for the high abundance of the *Aedes aegypti* vector. This unique characteristic of *Aedes aegypti* increases the potential for an outbreak should the virus be introduced into these areas (25).

One key finding in this study was the detection of *Aedes albopictus* in Madina. The presence of this vector has been documented before in Ghana, but this is the first time it has been recorded in such relatively high numbers (26). In a previous study, very few mosquitoes were identified as *Aedes albopictus* (Suzuki et al, 2016) whilst relatively higher numbers of the species were identified in this study. These mosquitoes are highly invasive species and numbers such as these could be an indication of a well-established population of *Aedes albopictus* mosquitoes locally. *Aedes albopictus* is another competent vector of DENV, CHKV and ZIKV and the presence of two competent vectors of these viruses in Madina put the community at an even greater risk for the transmission of these viruses (27–29). One important species identified in the forest area was *Aedes vittatus*, which is known to be a principally forest and savannah species of the *Aedes* mosquito. In some parts of Africa, *Aedes vittatus* has been linked to the spread of yellow fever and has also been experimentally proven to spread ZIKV (30–32).

Water holding containers in and around homes were inspected for the presence of the aquatic stages of the *Aedes* mosquito to allow for the estimation of the larval risk indices. Both study sites recorded high values for all indices except the House index for Madina which fell within the moderate risk range. These high values observed indicate a high risk of transmission should the Dengue, Zika and Chikungunya viruses be introduced into these communities through the presence of a positive human host, reservoir or vector. The relatively lower value observed for the House index in Madina could be attributed to the use of Polytanks^®^ (large water storage containers) and plastic barrels for storage of water in the community. These water storage containers were either empty or were properly covered; preventing the mosquitoes from gaining access to it as a potential breeding site. The community also has regular supply of water therefore there is little need for water storage.

Most containers found positive were observed in water holding containers outside of human dwellings such as discarded bicycle and car tyres, and cans. A high number of positive tyres was also recorded in the Achimota forest which were discarded near the forest office buildings. The study confirmed findings from other studies which showed that discarded tyres had a high positivity rate for *Aedes* larvae compared to other water holding containers. This may be because water collected in these tyres are left undisturbed for long periods of time. The tyres also provide a dark, cool and shady environment allowing the immature stages of the *Aedes* mosquito to grow in optimal conditions (33).

Adults and larvae were identified and pooled for detection of DENV, ZIKV and CHIKV. All pools analysed were found negative for the viruses of interest, is in line with other studies that have been conducted in other sites in Ghana (13,22). Some studies have suggested that *Aedes aegypti* mosquitoes originating from the African continent may be less competent vectors of these flaviviruses (34,35) because the ability of these mosquitoes to serve as competent vectors is influenced by several biotic and abiotic factors. Some of these viruses are extremely vector specific and some factors such as the bacterial communities associated with the vectors may interfere with the ability of the virus to replicate or be transmitted (36–38). Other studies have also indicated that Ghanaian strain of *Aedes aegypti* were refractory to infection by DENV serotype 2 compared to colonies from Vietnam (39).

Despite these reports, the *Aedes* mosquito population in Ghana are competent vectors of yellow fever virus and findings of this study reiterate the high risk they potentially pose to transmission other arboviruses. There are, however, several factors and avenues that still need to be studied to fully understand the population structure and vector competence of *Aedes*, and the role of primate reservoirs in arboviral transmission in Ghana.

## Conclusions

The study revealed that *Aedes aegypti* was the most abundant species of mosquitoes in the two study sites. Relatively high numbers of *Aedes albopictus* detected in Madina could signify the establishment of a local population. *Aedes* mosquitoes were also observed to be breeding close to human habitats. Although all pools were negative for DENV, ZIKV and CHKV, the high larval and ovitrap indices recorded indicate that there is a great potential for an outbreak of these viruses in both communities should they be introduced into the population through the presence of a positive human host, a zoonotic reservoir or an imported positive vector. The detection of *Aedes vittatus* also raises a cause for alarm since it has been implicated in the transmission of yellow fever and has been proven experimentally to transmit ZIKV.

It is also recommended that a national surveillance program be put in place to monitor the species abundance, diversity and risk of transmission of arboviral diseases in the country. The program should also periodically screen vectors for the presence of viral infections. Further studies should also be conducted in the country to test the vector competence of the *Aedes* species implicated in the transmission of arboviral diseases. This would help in the early detection of an outbreak and aid in the planning and implementation of effective and efficient control strategies.

## Acknowledgements

We are grateful to the staff of the Department of Parasitology and Vestergaard-NMIMR Vector laboratories especially Seraphim N.A Tetteh, Ibrahim K. Gyimah and Joseph Abraham for their support throughout the study. We also thank staff of the NMIMR Virology department, especially Eudocia Esinam Agbosu for her assistance in performing the viral detection assays. This study was supported by funding from BANGA Africa-Team Research Grant (UG-BA/TRG-006/2020-2021) to SKD. The funders had no role in the study design, data collection and analysis, decision to publish, or preparation of the manuscript.

## Author contributions

SKD, JA, NEA-B conceptualized the study. SKD, JA, KKF, SP-B, NEA-B, KB, MO discussed and designed the study. NEA-B, GKA, SSA, HAB, JHNO, RP, EAA, MO, conducted field sampling. NEA-B, GKA, SSA, DA-B, MO performed laboratory experiments. SKD, RQ supervised NEA-B studentship. NEA-B, JA, SKD analysed results and drafted the manuscript. All authors reviewed, edited, and approved the final manuscript.

